# Reference genomes for Australian *Cryptosporidium hominis* and *C. parvum* from humans

**DOI:** 10.64898/2026.11.26.769394

**Authors:** Marielle Babineau, Karolina Mercoulia, Michelle Sait, Sunita B. Sumanam, Taylor Harshegyi-Hand, Louise M. Judd, Norelle Sherry, Jake A. Lacey, Anson V. Koehler, Ryan Wick, Robin. B. Gasser, Benjamin P. Howden, Torsten Seemann

## Abstract

*Cryptosporidium hominis* and *C. parvum* are important protozoan pathogens of humans worldwide, and genomic surveillance is increasingly being used to investigate transmission and outbreaks. However, no complete *Cryptosporidium* genome is available from Australia, and many common *Cryptosporidium gp60* subtypes lack high-quality genome assemblies worldwide. This study generated the first Australian reference genomes for two clinically relevant, zoonotic *Cryptosporidium* species – *C. hominis* and *C. parvum* from human hosts, as well as a scaffold-level genome for Australian *C. meleagridis*. Oocysts were purified from eleven clinical faecal samples and whole-genome sequencing was performed using short-read technologies. Three samples belonging to different species were also sequenced using long-read technology, with assemblies generated, telomere recovery undertaken and genomes annotated. High-quality reference *C. hominis* IaA12R3 genome (8 T2T chromosomes, 9.2 Mb, 4069 genes, 98.4% completeness), near-complete *C. parvum* IIaA20G3R1 genome (8 chromosomes, 9.2 Mb, 4004 genes, 98.2% completeness), and a draft *C. meleagridis* IIIgA31 genome (113 scaffolds, 9.3 Mb, 4075 genes, 97.2% completeness) were generated and had high nucleotide similarity (98-99%), gene cluster orthology (99%) and genome synteny relative to available international reference genomes. Phylogenomic analyses with global *C. hominis* and *C. parvum* genomes were performed to characterise the relationships of Australian isolates globally showing Australian *C. hominis* and *C. parvum* forming well-supported clades with North American European isolates. Whole-genome analyses of eight Australian *C. hominis* isolates revealed markedly better resolution of genetic relationships than *gp60* subtyping alone. This study provides the resources for national genomic surveillance in Australia and expands the genomic representation of clinically relevant *Cryptosporidium* subtypes. Broader application of genomic surveillance has the potential to enhance outbreak investigation, improve understanding of transmission dynamics and support One Health approaches to cryptosporidiosis control.

**Impact Statement:** *Cryptosporidium* is a pathogen of significant public health, veterinary and environmental importance globally, causing a high burden of disease in children in low- and high-income countries. Genomic surveillance is emerging as an important tool to detect, control and prevent transmission of many pathogens, including *Cryptosporidium*. This approach requires high-quality local reference genomes to provide robust reconstruction of transmission clusters and SNP thresholds. This study presents the first Australian reference *Cryptosporidium* genomes from the two most common human-infective species in Australia, *C. hominis* and *C. parvum*, and representing *gp60* subtypes for which there were previously no genome assemblies available globally. The phylogenomic relationships between *Cryptosporidium* isolates from Australia and abroad show clustering with North American isolates. Using the new *C. hominis* reference genome, whole-genome analysis of eight Australian clinical *C. hominis* isolates demonstrates the readily applicable genomic resources presented here to implement genomic surveillance of *Cryptosporidium* in Australia.

**Data Summary:** All supporting data and protocols have been provided within the article or as supplementary data files. Detailed oocyst capture and DNA extraction protocol are provided in Supplementary Methods. The ONT and Illumina reads, annotated genome assemblies (n=3), and Illumina reads only (n = 8) from 11 isolates are available under PRJNA1465063.

## INTRODUCTION

Protozoan parasites of the genus *Cryptosporidium* cause cryptosporidiosis, a gastrointestinal disease that is often self-limiting but can be life-threatening in children and immunocompromised individuals [1]. *Cryptosporidium* is transmitted orally through contaminated water, food or direct transmission involving humans and/or animals [2]. Surveillance and prevention are the most important strategies due to the limited availability of effective drugs and the absence of a vaccine for use in humans and animals.

The genus *Cryptosporidium* currently comprises more than 45 species, 19 of which have been identified in humans [3]. *Cryptosporidium hominis* and *C. parvum* are the most common species found in humans worldwide, with *C. parvum* being the most common species in Europe, Canada and New Zealand, whereas *C. hominis* appears to be more common in Australia, the USA, South Asia and sub-Saharan Africa [2, 4–13]. Molecular surveillance of cryptosporidiosis has been implemented in the UK, Denmark, France and the USA [9, 14–16] using different approaches such as multi-locus sequence typing, 60 kDa glycoprotein (*gp60)* subtyping, and whole-genome sequencing. The genetic diversity within *Cryptosporidium* species is characterised by nucleotide variations within the *gp60* gene [17]. Routine molecular surveillance of cases in these countries has allowed for the rapid detection of local outbreak clusters, the linkage of outbreaks to international transmission events, the identification of zoonotic and waterborne sources and the improvement of disease control strategies [6, 12, 18–20].

To date, genome assemblies are available for 71 isolates representing 17 *Cryptosporidium* species. *Cryptosporidium* genomes have eight chromosomes with a genome size of 9.0 to 9.2 Mbp and approximately 4,000 protein-coding genes [21, 22]. The first complete *Cryptosporidium* genome was obtained from *C. parvum* in 2004 and, following revisions, has revealed complex subtelomeric regions [23–25]. No public genomes are available for the Australian continent [21, 26]. Despite the 264 *gp60* subtypes identified within *C. parvum* and *C. hominis* globally, only 20 have genome assemblies available: seven *C. hominis* subtypes and 13 *C. parvum* subtypes [21, 27]. High inter- and intra-specific genomic diversity has been observed, as well as signatures of adaptation to local selection pressures, indicating genomic diversity associated with geographic isolation and local adaptation [12, 28–30]. These observations suggest that Australian *Cryptosporidium* isolates may be distinctive phylogenomically. Unique Australian *Cryptosporidium* species, genotypes and subtypes have been identified using *gp60* [4, 31–37]. Geographically, it might be expected that Australian *Cryptosporidium* spp. would be more closely related to South Asian and New Zealand isolates; however, evidence of long distance dispersal of *C. parvum* [19, 38] suggests that Australians isolates could be genetically more similar to European or North American isolates. Given the high incidence of clinical cryptosporidiosis in Australia compared to other high-income countries, and that the majority of cases appear to be acquired locally in Australia [39, 40], there is a need for national genomic resources to enable surveillance [41]. However, clinically relevant, high-quality, Australian *Cryptosporidium* genomes are needed to enable molecular surveillance.

This study aimed to generate high-quality complete *C. hominis*, *C. parvum* and *C. meleagridis* genomes from Australian clinical samples, place them in the global *Cryptosporidium* population structure, and use the newly generated reference genome to assess phylogenetic relationships among closely related Australian *C. hominis* isolates using whole-genome sequencing.

## METHODS

### Clinical samples selection and oocyst purification

In 2024 and 2025, clinical faecal samples were referred to the Microbiological Diagnostics Unit Public Health Laboratory (MDU PHL) from local diagnostic service providers for testing and biobanking at 4 °C. Based on previous *gp60* subtyping [4], one sample from each of the three most commonly reported species found in Australians, *C. hominis, C. parvum* and *C. meleagridis* were selected for complete genome assembly (Table 1). In addition, a subset of eight *C. hominis* clinical samples collected in 2024 representing different *gp60* subtypes [4] was selected for shotgun whole-genome sequencing (Table 1). Oocysts were purified from 1 g of fresh faeces using the CryptoBeads ® kit (BioPoint Pty, Sydney, Australia) using a modified protocol (Supplementary Methods). Purified oocysts were enumerated with a haemocytometer (Hausser Bright Line, Stemcell Technologies, USA) using a light microscope (100-times magnification) and preserved in 500 µl of DNA/RNA Shield solution (Zymo Research, Australia).

**Table 1.** Genome sequencing and assembly status of 11 Australian *Cryptosporidium* spp. samples from human hosts.gp60; 60 kDa glycoprotein gene. Paired end (PE) reads number post quality filtering.

| Isolate | Species | *gp60* subtype (NCBI Accession number) | Sequencing type (depth) | Genome assembly status (number) | NCBI Accession number (assembly) |
|---|---|---|---|---|---|
| **AUSMDU00120773** | *C. hominis* | IaA12R3 (NA) | ONT (358x) + Illumina (59x) | Chromosomes (8) | SAMN59746842 (GCA_058640825.1) |
| **AUSMDU00108975** | *C. parvum* | IIaA20G3R1 (PX092536) | ONT (258x) + Illumina (200x) | Chromosomes (8) | SAMN59746841 (GCA_059910425.1) |
| **AUSMDU00108966** | *C. meleagridis* | IIIgA31 (PX092399) | ONT (157x) + Illumina (167x) | Scaffold (113) | SAMN59746839 (GCA_059910485.1) |
| **AUSMDU00108930** | *C. hominis* | IfA12G1R5 (PX092510) | Illumina (164x) | PE reads (10,784,302) | SAMN59746835 |
| **AUSMDU00108967** | *C. hominis* | IaA16R3 (PX092615) | Illumina (227x) | PE reads (15,365,456) | SAMN59746840 |
| **AUSMDU00108955** | *C. hominis* | IdA15G1/IbA9G3 (NA) | Illumina (105x) | PE reads (7,054,764) | SAMN59746838 |
| **AUSMDU00101309** | *C. hominis* | IaA12R3 (NA) | Illumina (157x) | PE reads (10,100,554) | SAMN59746834 |
| **AUSMDU00108949** | *C. hominis* | IaA16R4 (PX092616) | Illumina (164x) | PE reads (10,510,494) | SAMN59746836 |
| **AUSMDU00108953** | *C. hominis* | IdA15G1 (PX092549) | Illumina (120x) | PE reads (8,300,362) | SAMN59746837 |
| **AUSMDU00101221** | *C. hominis* | IaA12R3 (PX092560) | Illumina (167x) | PE reads (10,681,916) | SAMN59746833 |
| **AUSMDU00108968** | *C. hominis* | IaA24R1 (PX092557) | Illumina (141x) | PE reads (9,816,628) | SAMN60747431 |

### DNA extraction and amplification

Preserved oocysts were diluted in purified water and centrifuged at 5,000 g for 7 min to precipitate oocysts, the supernatant was discarded and the precipitated oocysts were resuspended in 500 µl of DNA Lysis Buffer (Supplementary Methods). DNA was extracted using a modified phenol-chloroform protocol (Supplementary Method). DNA quality and quantity were assessed using the 4200 Tapestation system (Agilent Technologies, Australia). If the DNA quantification was below 1 ng/µl, DNA was amplified using the Repli-g Mini kit (Qiagen, Australia) following the manufacturer’s protocol. Amplified DNA was debranched using endonuclease I (New England Biolabs, UK) using a modified protocol (Supplementary Methods). With the exception of AUSMDU00120773, all isolates underwent whole genome amplification prior to short and long read sequencing.

### Short read sequencing

All 11 samples were subject to short reads WGS. Sequencing libraries were prepared with Illumina DNA-Prep (quarter reagents) and sequencing was performed on the Illumina NextSeq 2000 platform with 2 x 150 bp paired-end reads. Resulting reads underwent adapter removal, trimming and quality trimming using fastp v0.23.2 [42] with default parameters and a minimum length of 36. The median sequencing depth was 161.

### Long read sequencing

Long reads sequencing was performed on one isolates from each species of interest based on previous speciation, subtyping and fresh material availability: AUSMDU00120773, AUSMDU00108975 and AUSMDU00108966 (Table 1). Between 43 and 168 ng of DNA (non-amplified AUSMDU00120773 DNA and amplified DNA for the remaining isolates) was used for Oxford Nanopore Technologies (ONT) sequencing on the PromethION P2 Solo (FLO-PRO114M). DNA libraries were prepared using the Ligation sequencing gDNA-Native Barcoding kit 24 V14 (SQK-NBD114.24) following the manufacturer’s instructions. The super-high accuracy base-calling option (dna_r10.4.1_e8.2_400bps_sup@v4.3.0) and sequencing was performed for 72 h. Reads below a mean quality score of 10 were eliminated using fastplong v0.7.1 [42]. After fastplong, reads were passed through Filtlong v0.2.1 (https://github.com/rrwick/Filtlong) with a minimum length of 1,000 bp and retaining 95% of reads.

### Genome assemblies and annotation

Long-read assemblies were generated using Flye v2.9.6 [43] specifying a genome size of 9 Mb (-- genome-size 9m) using the command --*nano-hq*; the option *--no-alt-contig* was used for *C. meleagridis* to remove a high number of highly similar duplicated contigs differentiated by only 1-2 SNPs assessed as alternative alleles (see Results). The resulting contigs were compared against all nucleotide sequences in GenBank using BLASTN v2.13.0 against the standard core nucleotide database (core_nt) to eliminate contigs yielding a high nucleotide similarity (>90%) to organisms other than *Cryptosporidium* spp. Draft assemblies were polished using Medaka v2.1.1 (github.com/nanoporetech/medaka) using default parameters. The short read data were used to correct errors, especially in repeat regions, in the long read assembly using Polypolish (https://github.com/rrwick/Polypolish) with default parameters.

Telomeric regions were recovered and extended using Teloclip v0.3.2 (github.com/Adamtaranto/teloclip) using long reads data. Telomere recovery was assessed using tidk v0.3.2 [44] with a minimum of 20 TTAGG repeats [45] required with the last 1,000 bp of the 3’ and 5’ ends of each assembled contig to indicate successful telomere recovery. Each assembled contig was aligned against a reference genome assembly corresponding to the same species – *C. parvum* CpBGF ASM3523276v1 (GCA_035232765.1 [25]) and CpBCM-BGF ASM1984411v2 (GCA_019844115.2) [46], *C. hominis* UdeA01 (GCA_002223825.1 [47]) and *C. meleagridis* TU1867 (GCA_039657295.1 [48]) – using BLASTN to match chromosome homology and strand orientation, and visualised in kablammo (kablammo.wasmuthlab.org). Contigs that did not align against their specified reference genome and showed a high nucleotide identity (e-value < 1e^-3^ across 90% of the sequence length) to organisms other than *Cryptosporidium* spp. were eliminated and classified as contamination. Assembly statistics was obtained using QUAST v5.3.0 [49]. Assembly completeness was assessed using BUSCO v5.4.5 [50] with the coccidia_odb10 database with default parameters. Synteny was assessed by identifying blocks of locally collinear nucleotides in ProgressiveMauve with default parameters [51] using the CpBGF and CpBCM-BGF as reference for *C. parvum* and the UdeA01 for *C. hominis*. Pairwise nucleotide and SNP differences between newly generated and reference genome assemblies were calculated using the dnadiff command and visualised using the delta-filter (-i 98 -l 1000) and mummerplot commands in Mummer4 v4.0.1 [52]. For *C. parvum*, the chromosomes of the newly generated genome were ordered and orientated based on the reference assembly with the highest average pairwise identity.

Genome annotation was performed using Companion [53] using default parameters with reference genomes *C. parvum* Cp IOWA II (ASM16534v1, GCF_000165345.1 [23]), *C. hominis* TU502_2012 (GCA_001593465.1), and *C. meleagridis* UKMEL1 (GCA_001593445.1). The *C. hominis* UdeA01, *C. parvum* CpBGF, *C. parvum* CpBCM-BGF and *C. meleagridis* TU1867 are not currently available in Companion. The assignment of protein-coding gene clusters between the Australian assemblies and selected reference genomes was visualised using OrthoFinder v2.5.4 [54] based on Diamond hits (e-value cutoff of 10^-3^) of the translated protein sequences. Visualisation of telomere repeats, GC content, chromosome similarity, gene density and gene coverage was performed using shinyCircos [55].

The subtype of all sequenced isolates was assessed by blasting (blastn) known subtype nucleotide sequences of the single-copy gene *gp60* against the genomic assemblies, retrieving the *gp60* sequence and assessing subtype following most recent nomenclature [17]. Quality filtered (fastp) paired reads from isolates which were only sequenced using short-read technology were assembled *de novo* using SPAdes v4.0.0 (ablab.github.io/spades). The genomic assemblies *gp60* sequences were compared to *gp60* subtype previously obtained by PCR and Sanger sequencing in 2024 when available [4].

### Phylogenomic analysis

All available SRA run ID and associated metadata for short-read genomic sequences for *C. hominis* and *C. parvum* were retrieved from NCBI on 30/06/2026 using the following filters: Assay type: wgs and wga, Library source: genomic, library layout: paired, platform: Illumina. The metadata was filtered manually to remove SRA runs generated from experimental laboratory-derived strains unsuitable for phylogenetic comparisons. Fastq files were retrieved using sracha v0.5.0 (github.com/rnabioco/sracha-rs). Speciation was assessed based on read taxonomic assignment using kraken2 v2.17.1 with the PlusPF database. Quality assessment and speciation was performed using bohra v3.7.2 (github.com/MDU-PHL/bohra) using the basic pipeline. Runs with less than 20 Mbp, less than 40x depth, showing a speciation match (match #1) to a species other than *Cryptosporidium*, and runs showing more than 40% of reads with unassigned speciation were eliminated (Supplementary Table 1). Pairwise SNP differences were calculated using the bohra comparative pipeline with *C. parvum* CpBGF (GCA_035232765.1) and *C. hominis* UdeA01 (GCA_002223825.1) as reference genomes using snippy v4.4.5 to assess SNP variants (fraction of reads threshold of 90% with a minimum of 10 reads, --minfrac 0.9, --min cov 10) and using a soft core allowing for 5% of missing taxa for each SNP (fuzzy-core-prop 0.95). Ambiguous sites were excluded from the analysis unless they were only present in 5% or less of the isolates evaluated. The alignment of core polymorphic SNPs (core_filt.aln) was used to generate a maximum likelihood phylogenetic tree using the GTR+I+G nucleotide substitution model using IQ-TREE v3.1.2 with 1000 ultrafast (-B) bootstrap replicates. Phylogenetic trees were modified (tree scale adjusted to discrete SNP number) and visualised using the R package ggtree [56]. Lineage and subspecies assignment for *C. hominis* and *C. parvum* followed previously published taxonomic classifications [19, 30, 57, 58]. Available *gp60* subtype provided as metadata from NCBI was used to annotate the trees, when not provided, the *gp60* sequence was retrieved from the short-read R1 and R2.fastq using an in-house script which used the cryptogenotyper [59] reference *gp60_ref.fasta* as query database, the retrieved sequence was then used as input in cryptogenotyper to confidently identify subtype. Pairwise nucleotide and SNP differences between genome assemblies were calculated using the dnadiff as above.

Genomic diversity within Australian *C. hominis* isolates was assessed using the same phylogenomic method as above, with the exception that AUSMDU00120773 genome assembly was selected as the reference. The whole-genome SNP tree was compared to a *gp60* neighbour-joining tree with 1000 bootstrap replicates using a MUSCLE alignment of the previously published sequences in NCBI (Table 1). The AUSMDU00120773 *gp60* sequences, as well as any isolate showing discordance between WGS and previously published *gp60*, was retrieved from their genome assembly following the above methods.

## RESULTS

### Australian reference genomes

The genomes of Australian *C. hominis* and *C. parvum* were assembled into eight chromosomes (Table 1). The *C. hominis* assembly was a complete T2T genome, including 16 telomeres with a genome size of 9.2 Mb comprising 4,069 genes and with an inferred 98.4% single-copy gene recovery (Table 2).

**Table 2.** Genome assembly and annotation statistics for the Australian *C. hominis* and *C. parvum* reference genomes (AUSMDU) compared with the *C. hominis* UdeA01 and *C. parvum* CpBGF and CpBCM-BGF respective. Assembly CpBCM-BGF is not annotated.

|  | ***C. hominis*** |  | ***C. parvum*** |  |  |
|---|---|---|---|---|---|
| **Isolate** | AUSMDU00120773 | UdeA01 | AUSMDU00108975 | CpBGF | CpBCM-BGF |
| **Number of assembled chromosomes** | 8 | 8 | 8 | 8 | 8 |
| **Number of contigs** | 8 | 12 | 8 | 8 | 8 |
| **Number of telomeres** | 16 | 9 | 8 | 16 | 13 |
| **Genome size** | 9,209,345 | 9,068,851 | 9,175,495 | 9,259,183 | 9,197,619 |
| **Genome coverage** | 358x | 68x | 258x | 2000x | 53x |
| **Longest contig (bp)** | 1,359,803 | 1,322,947 | 1,354,268 | 1,379,419 | 1,355,650 |
| **N50** | 1,106,661 | 1,103,974 | 1,109,704 | 1,107,426 | 1,108,772 |
| **N90** | 991,162 | 988,320 | 994,832 | 992,060 | 993,129 |
| **L50** | 4 | 4 | 4 | 4 | 4 |
| **Number of N per 100 kbp** | 0 | 93.73 | 0 | 0 | 0 |
| **Overall GC %** | 30.0 | 30.1 | 30.1 | 30.0 | 30.1 |
| **Number of genes** | 4,069 | 3,819 | 4,004 | 4,841 | NA |
| **Protein-coding genes / non-coding genes** | 3,974 / 95 | 3,817 / 2 | 3,921 / 83 | 3,928 / 913 | NA / NA |
| **Complete & single-copy gene** (%, M; Missing, F; Fragmented) | 494 (98.4%, 5M, 3F) | 494 (98.4%, 5M, 3F) | 493 (98.2%, 5M, 4F) | 493 (98.2%, 5M, 4F) | 494 (98.4%, 5M, 3F) |

This complete T2T *C. hominis* genome is similar to the only other complete T2T *Cryptosporidium* genome available, the *C. parvum* CpBGF assembly in regards to genome size, longest contig and number of protein-coding genes [25]. Compared to the UdeA01 chromosome assembly, the Australian *C. hominis* genome represents a higher completeness assembly with all telomeres recovered, zero Ns per 100 kbp and a higher number of genes. The newly generated *C. parvum* genome recovered eight telomeres with a total genome size closer to CpBCM-BGF, and a number of protein-coding genes and single-copy gene recovery (493, 98.2%), highly similar to CpBGF (Table 2). The AUSMDU00108975 assembly showed a higher average nucleotide similarity to CpBCM-BGF (99.94%, 1,982 SNPs) compared to the CpBGF assembly (99.92%, 2,877 SNPs).

The orthologous protein-coding gene clusters for *C. hominis* was 98.9% and 99.8% for *C. parvum* using TU502_2012 and Cp IOWA II as annotation reference respectively (Figure 1). Between 10 and 45 genes were identified in the Australian *C. parvum* and *C. hominis* assemblies respectively, which were not present in their respective annotation reference. Between 77% to 100% of these unshared genes were identified as conserved hypothetical proteins, limiting any interpretation of functional differences (Supplementary Table 2). However, four putative transferase genes (glutamine amido-transferase and ubiquitin transferase) were identified within Australian *C. parvum*, while fifteen unshared protein-coding genes were identified from the Australian *C. hominis*; seven linked to DNA/RNA metabolism, four related to transport and signalling, and two to genes related to mitochondrial metabolism (Supplementary Table 2). The two latter genes likely relate to the functionally reduced mitosome in *C. hominis*, a remnant of a largely lost mitochondrial genome [60]. There were no unshared genes clusters in common between the Australian *C. hominis* and *C. parvum*.

**Figure 1.**
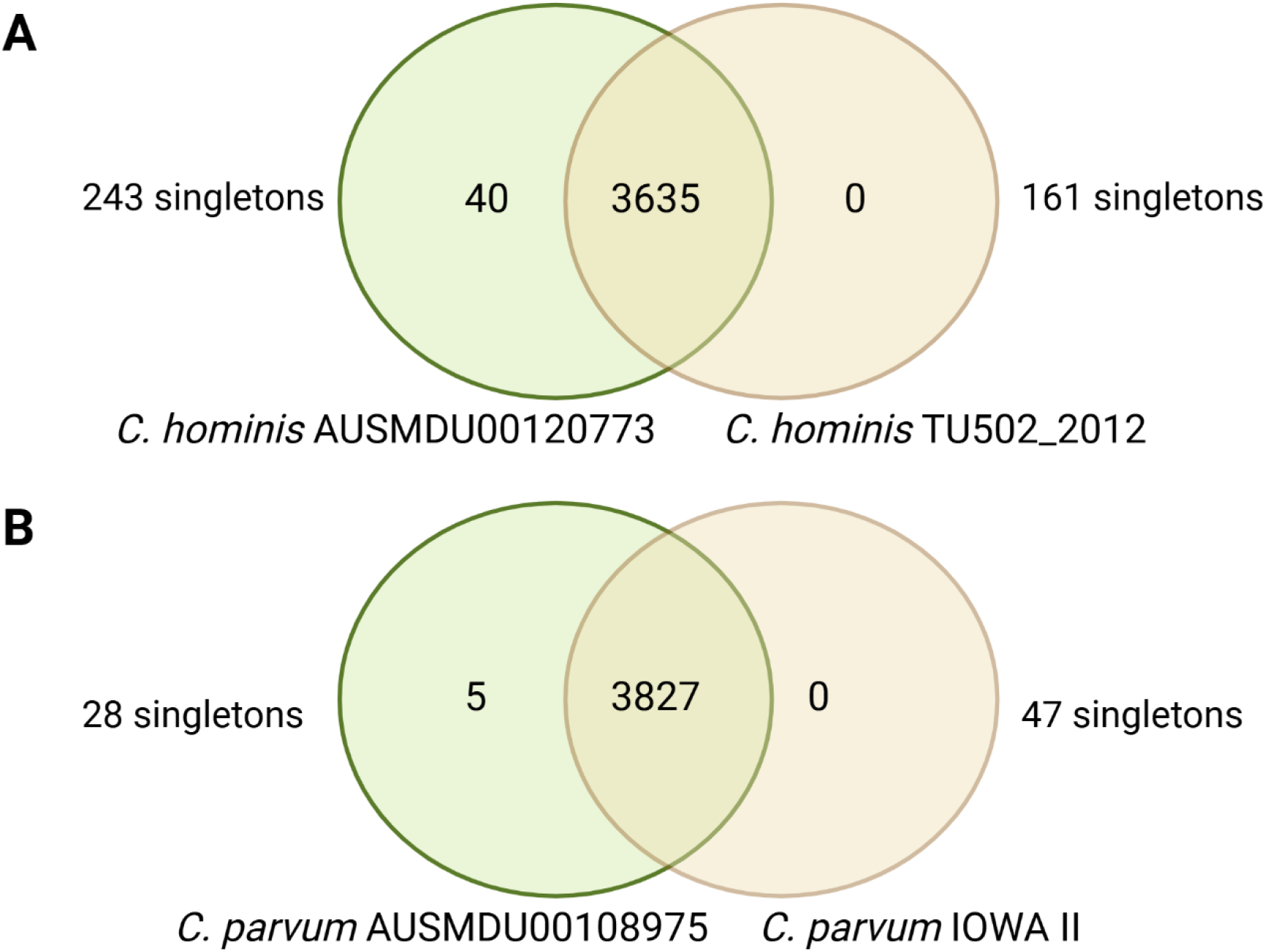
Orthologue protein-coding genes clusters for *C. hominis* AUSMDU00120773 (A) and *C. parvum* AUSMDU00108975 (B). Shared gene clusters based using a minimum of two genes per cluster. Singletons are the number of genes not assigned to any clusters.

The two Australian chromosome-level genomes showed chromosome 8 as the largest and chromosome 1 as the shortest (Figure 2), as observed previously [23, 46, 47]. The GC content and gene coverage across the genomes were consistent with decreases at the chromosome ends. For both isolates, gene density was highest in chromosome 2. Almost all regions of the genomes of the Australian isolates were similar to those in respective reference genomes. The Australian *C. hominis* genome showed a higher proportion of nucleotide divergence relative to the UdeA01 genome than was observed for *C. parvum* relative to CpBCM-BGF due to the more distant relationship between the *C. hominis* genomes representing different *gp60* subfamilies (Ia vs Ie) compared to the *C. parvum* genomes (all IIa subfamily). Nucleotide similarity of chromosome 2 between the *C. parvum* AUSMDU00108975 genome and CpBGF was lower compared to CpBCM-BGF (Figure 2B) with an average blastn bitscore of 364,160 vs 914,000 respectively supporting the higher genome similarity of the newly generated assembly and CpBCM-BGF.

**Figure 2.**
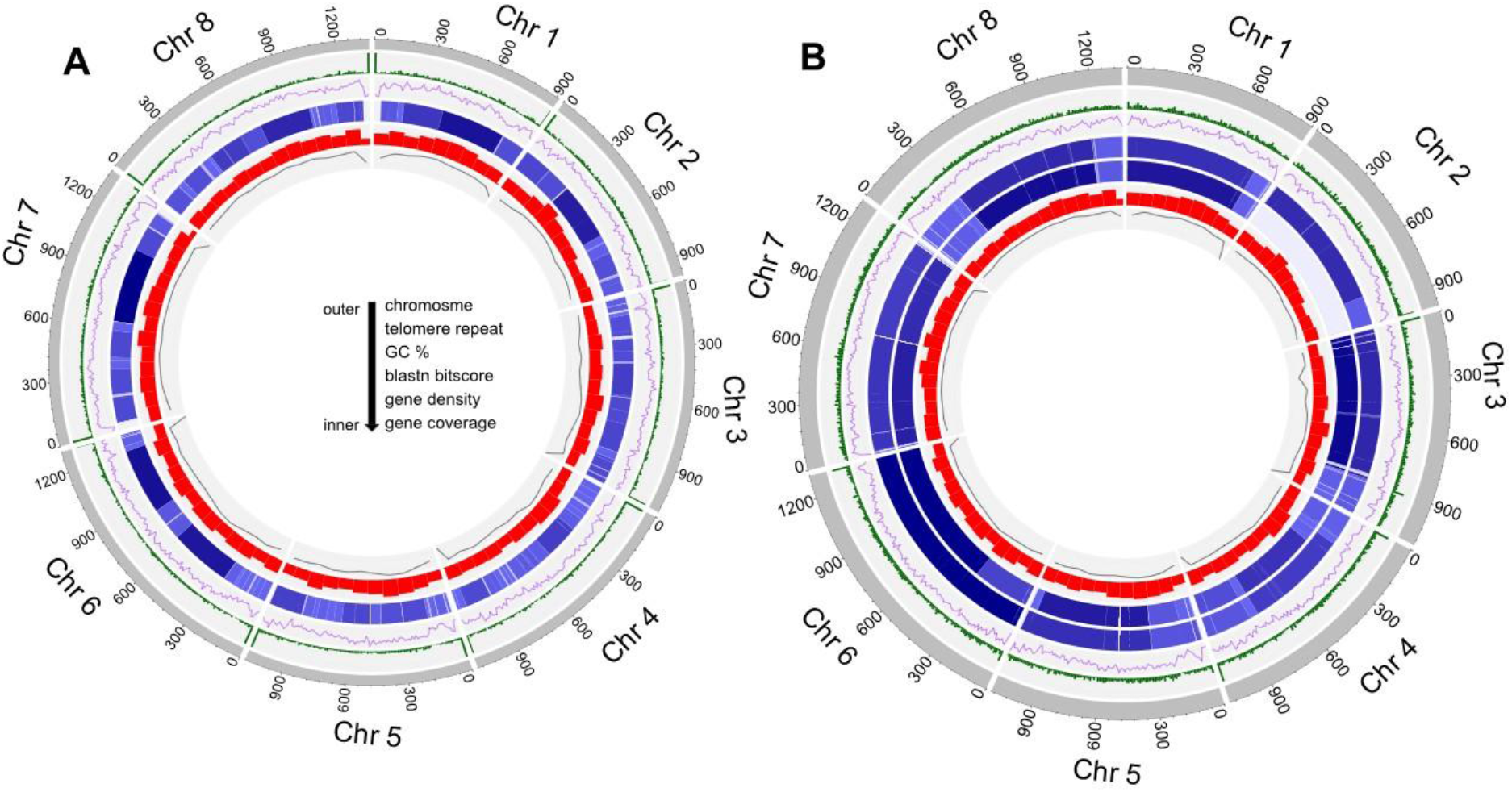
Genome assembly data and nucleotide similarity for *Cryptosporidium hominis* isolate AUSMDU00120773 (A) and *Cryptosporidium parvum* isolate AUSMDU00108975 (B). Circle plot rings (outer to inner): chromosome size (KB), number of telomere repeat TTAGG/CCTAA per 10,000 bp, GC percentage per 10,000 bp, bitscore blastn nucleotide similarity with the UdeA01 genome assembly (for *C. hominis*) and CpBCM-BGF and CpBGF (for *C. parvum;* layer 4 and 5 respectively) with light blue (>50) to dark blue gradient (>10,000), gene density per 100,000 bp, gene coverage per 100,000 bp.

There was syntenic variation at the nucleotide level between the Australian *C. hominis* genome and the UdeA01 genome (Supplementary Figure 1), with large inversions at both ends (137,938 bp at the 3’ end, and 385,062 bp at the 5’ end) of chromosome 4 as previously observed in *C. hominis* and *C. parvum* genomes [24, 61]. There was no major syntenic variability between the Australian *C. parvum* and the CpBCM-BGF and CpBGF which were highly collinear (Supplementary Figure 2). The highly similar assembly statistics, gene recovery, orthology and synteny observed supports the genomic similarity and completeness of the Australian *C. hominis* and *C. parvum*.

The *C. meleagridis* genome was assembled into 113 scaffolds (Supplementary Table 3). No telomeres were recovered and the genome showed a 2% greater size than the TU1867 genome while recovering a lower percentage of single-copy genes. The 113 scaffolds covered approximately 90% of the TU1867 chromosomes (data not shown); however, there was a high number of overlapping regions between scaffolds, causing the greater genome size and reflecting conflicting SNPs and indels, suggesting intra-sample genetic heterogeneity that could not be detected previously using Sanger sequencing of the *SSU* and *gp60* loci. Despite this limitation, the *C. meleagridis* genome showed a 99.6% recovery (3,607/3,619) of orthologous protein-coding gene clusters compared to UKMEL1 (Supplementary Figure 3), in line with the recovery observed for *C. hominis* and *C. parvum*. A single *gp60* subtype IIIgA31 (100% similarity to PX092399) was identified from the Australian *C. meleagridis* genome assembly consistent with previous Sanger sequencing, the multiple copies of *SSU* recovered were also 100% identical to the previously published sequence (PV981446) supporting genomic heterogeneity as opposed to mixed population within this isolate.

The *gp60* subtype of the Australian *C. parvum* genomic assembly was the same as previously determined based on Sanger sequencing, as well as for six of the nine *C. hominis* isolates. Isolates AUSMDU00101309 and AUSMDU00120773 were not previously subtyped. Isolate AUSMDU00108955, previously subtyped as IbA8G4 (PX092403), showed two different subtypes with subtype IbA9G3 (98% similarity with 17x coverage) and IdA15G1 (98% similarity with 74x coverage) each being retrieved from two different contigs within the short-read assembly.

### Global phylogenomic relationships of Australian isolates

A total of 321 short read *C. hominis* genomes were retrieved from NCBI including the nine Australian *C. hominis* isolates from this study, of which 303 passed quality control threshold. The most common exclusion criteria were depth below 40x (44%) and speciation to non-*Cryptosporidium* taxa (44%). An additional 20 isolates were excluded during the SNP calling due to low quality mapping to the reference and/or core SNPs not supported (Supplementary Table 1). The soft core of 244,343 SNPs alignment included 283 genomes including eight of the nine newly generated Australian *C. hominis* (Figure 3). Isolate AUSMDU00108930 was excluded from the SNP calling and core SNP tree reconstruction due to the low percentage of reads alignments to the reference genome (67% vs 97% in other isolates) and high percentage of reads speciating to human (match 2: *Homo sapiens* 17% vs 1-2% in other Australian isolates). The eight remaining Australian *C. hominis* isolates clustered in the *C. hominis hominis* subspecies and formed six well supported clades often with isolates from North America (four clades; all isolates from the USA) but also with European (e.g. UK), Asian (e.g. China) and African (e.g. Tanzania) isolates (three clades) all collected from human hosts. The AUSMDU00108949 isolate subtype IaA16R4 was sister to a clade containing isolates of the If subfamily but was separated by a high number of SNPs compared to other clades (Figure 3). Isolates belonging to the Ia subfamily, Australian and worldwide, were spread out in multiple well supported clades supporting previous evidence that the *gp60* Ia subtypes represent a large genomic diversity and are not monophyletic [12, 28]. The AUSMDU00108953 IdA15G1 isolate was closely related to USA isolate 46117 IeA11G3T3, the AUSMDU00108967 IaA16R3 was related to the isolate UKH30 IbA9G3, while AUSMDU00108968 IaA24R1 was closely related to two USA isolates (45690, 43739) both of subtype IaA16R3. The isolate AUSMDU00108955 of subtype IdA15/IbA9 formed a highly supported clade with African isolates (Afr18, Afr24, Afr15) of subtypes IdA15 and IdA17. The three IaA12R3 Australian isolates were most closely related to each other and were sister to a North American clade containing isolates of subtype IaA28 (30976, 43434). The *gp60* subtype of *C. hominis* isolates was not an accurate indication of the whole genome SNPs clustering based on this global dataset as previously reported [28]. The Australian *C. hominis* reference genome assembly AUSMDU00120773 was 99.90% similar to its sister-clade genome assembly of US isolate 30976 isolate (ASM148351v1; 995 core SNPs), supporting a close proximity to clinical USA isolates. The AUSMDU00120773 assembly was 99.91% similar to the distantly related assembly of isolate UKH4 IaA14 (ASM130532v1; 1,358 core SNPs) placed in the second earliest diverging clade within *C. hominis hominis* (Figure 3). The high and almost equal pairwise similarity percentage, despite different numbers of pairwise SNPs, suggests that large portions of the genomes are conserved between AUSMDU00120773, 30976 and with UKH4 with localised high-density SNPs areas, as corroborated by the alignment mummer plots.

**Figure 3.**
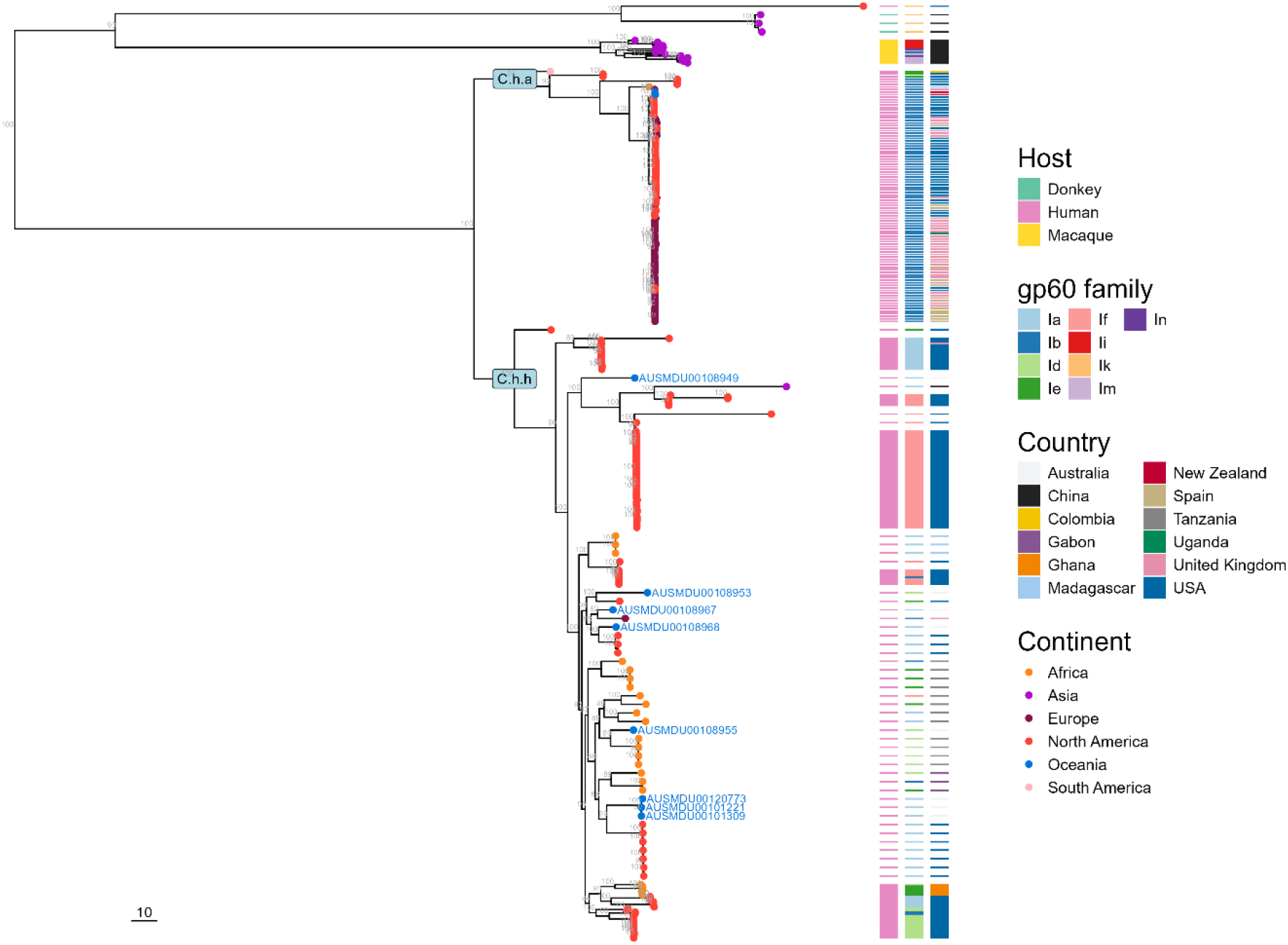
Maximum Likelihood phylogenomic tree based on 244,343 whole genome polymorphic SNPs from 283 *C. hominis* genomes. Bootstrap values above 80 shown above branches. Midpoint rooting used. Subspecies C.h.a; *C. hominis aquapotensis*, C.h.h; *C. hominis hominis*. Newly generated Australian isolates (AUSMDU) identified in blue. Tree scale bar set at 10 SNPs.

A total of 752 short read *C. parvum* genomes were retrieved from NCBI including the newly generated Australian *C. parvum*, of which 458 passed quality control criteria. The most common exclusion criteria was experimental laboratory-derived data (70%) followed by depth below 40x and non-*Cryptosporidium* speciation match (9% each), with an additional 16 isolates eliminated during the SNP calling analysis. The soft core of 250,020 SNPs alignment included 442 genomes (Figure 4). The Australian *C. parvum* belonged to subspecies *C. parvum parvum*, nested within Population 3 of the Western Lineage [19]. AUSMDU00108975 was in a highly supported clade with US isolate 42886 of subtype IIaA18G3R1. This clade, along with its sister clade containing two European isolates (Prt1, Spain1), were at the junction between a large European cluster and a large North American clade. The Australian *C. parvum* genome assembly was 99.94% similar to both the UKP4 genome assembly (ASM130541v1; 953 core SNPs) and the CpBCM-BGF (bottom of North American Pop3 clade) genome assembly (ASM1984411v2; 1,982 core SNPs), supporting the Australian isolate within the Western lineage and supporting an intermediate similarity between European and North American isolates.

**Figure 4.**
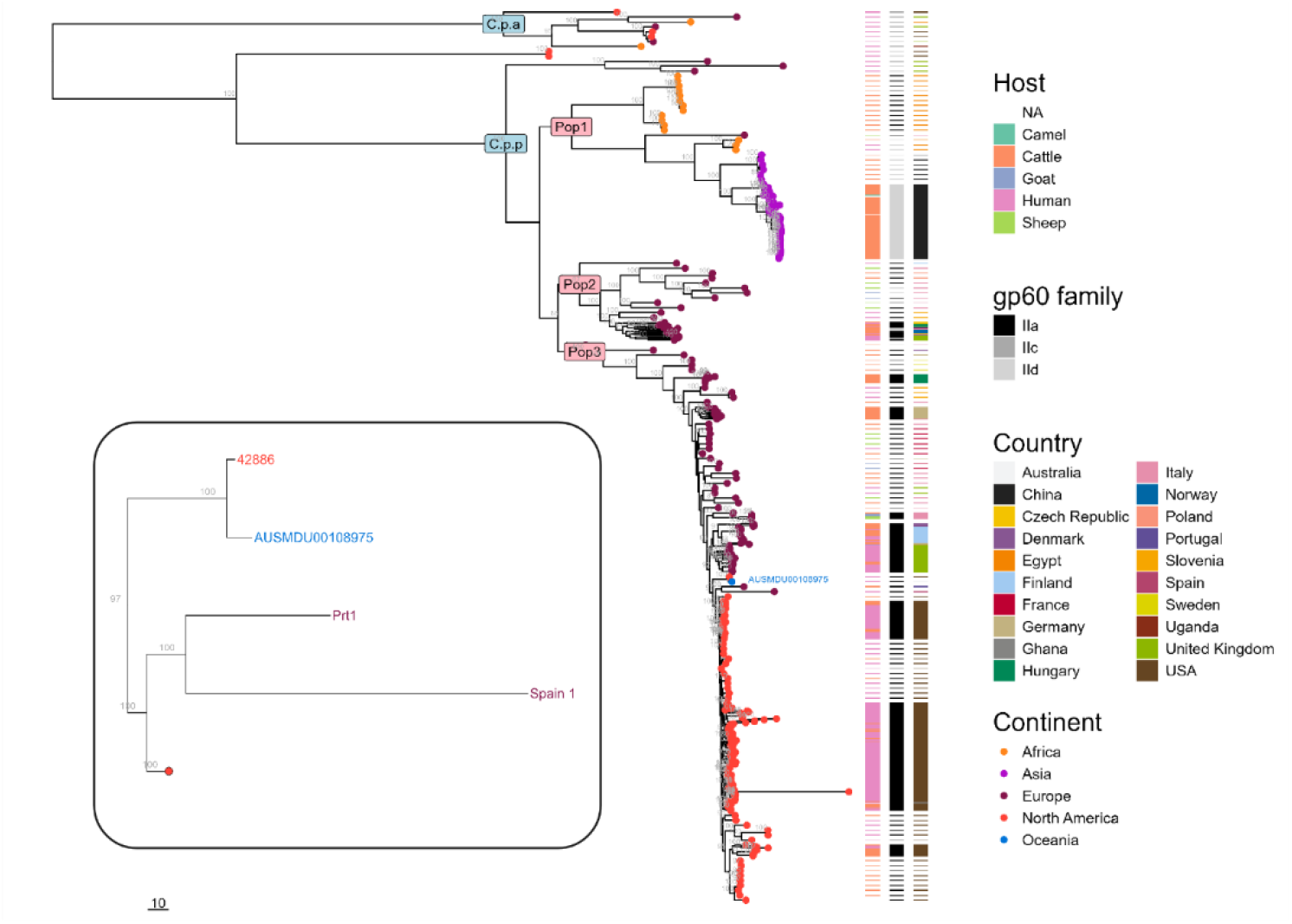
Maximum Likelihood phylogenomic tree based on 250,020 whole genome polymorphic SNPs from 442 *C. parvum* genomes. Bootstrap values above 80 shown above branches. Midpoint rooting used. Subspecies C.p.a; *C. parvum anthroponosum*, C.p.p; *C. parvum parvum*. Populations as Pop1; China/Egypt Lineage, Pop2; basal Western Lineage. Tree scale bar set at 10 SNPs.

### High genomic diversity of Australian *C. hominis*

The whole genome phylogenetic tree based on 2,896 soft core SNPs among eight of the nine Australian *C. hominis* isolates sequenced showed a more detailed resolution of genetic relationships between the Australian *C. hominis* clinical isolates compared to the *gp60* tree (Figure 5). Isolate AUSMDU00108930 was excluded for the same reasons as above. Contrary to the single locus tree, the whole genome SNPs (wgSNPs) tree showed a greater number of bifurcating clades with higher support values indicating a greater number of derived SNPs for each clade and taxa. The *gp60* and wgSNP trees showed some congruent relationships among isolates. In both analyses, the three IaA12R3 were highly similar, the AUSMDU0010867 and AUSMDU0010868 (both Ia subfamily) were shown to be closely related, and the Id isolates were sister to most of the Ia isolates. This finding supports the use of *gp60* subtyping as a moderately accurate, low-cost method to classify isolates at lower resolution. However, the wgSNPs tree displayed differences in the phylogenetic relationships; the six Ia isolates clustered in three different clades, two of those clades well supported. The position of heterogeneous isolate AUSMDU00108955 as more closely related to the IaA12R3 clade in the wgSNPs tree is further supported by the greater nucleotide similarity between this isolate and AUSMDU00101309 (99.95% identity, 1,041 GSNPs) compared to AUSMDU00108953 (99.93% identity, 1,405 GSNPs), however the position of AUSMDU00108955 had a low support value of 74 suggesting ambiguous position. The analysis without AUSMDU00108955 showed the same topology and relationships between other isolates but with higher bootstrap support for the position of AUSMDU00108953 (Supplementary Figure 4) again suggesting the position of isolate AUSMDU00108955 as ambiguous supporting a mixed population isolate. The Australian wgSNPs tree (Figure 5B) showed relationship highly congruent with the global wgSNPs *C. hominis* tree (Figure 3) which used different reference genomes. This finding is consistent with previous reports of discordance between *gp60*-based and whole-genome phylogenies [8, 28, 62].

**Figure 5.**
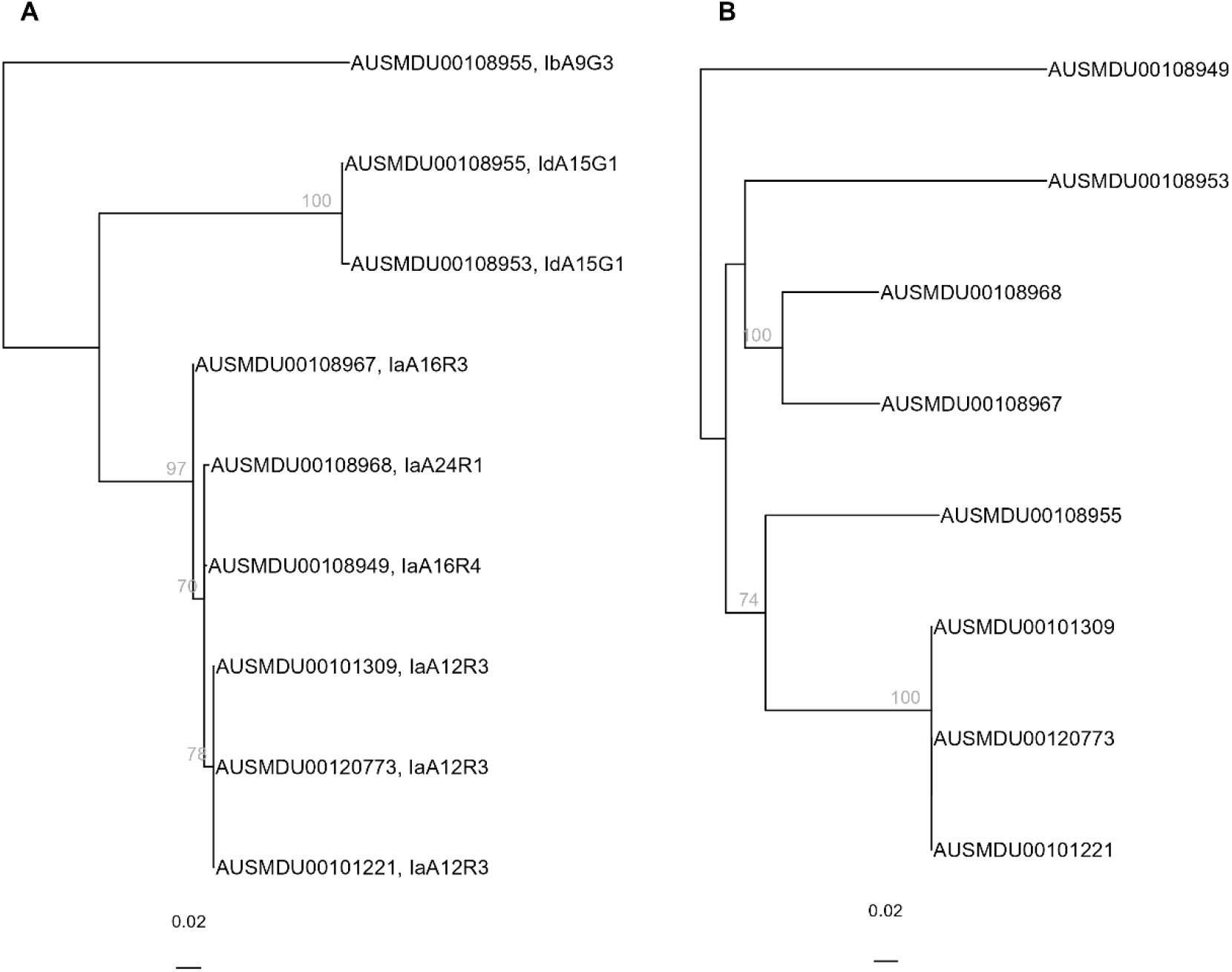
Phylogenetic relationships of eight Australian *C. hominis* isolates based on Neighbour-Joining reconstruction of *gp60* gene nucleotide alignment (A) and Maximum Likelihood reconstruction of 2,896 whole-genome short-read SNPs distances with isolate AUSMDU00120773 used as reference genome (B). Bootstrap support values shown above branches. Midpoint rooting used for both trees.

## DISCUSSION

This study provides the first three Australian *Cryptosporidium* genomes representing *gp60* subtype for which no genomes assemblies were previously available. The high levels of nucleotide similarity, orthology and synteny observed relative to other reference genomes further confirm that genomes from human-infective isolates are often very similar and syntenic [21]. The Australian *C. hominis* and *C. parvum* reference genomes contained a small number of unshared genes compared to their annotation reference, most of which were uncharacterised, a limitation also found previously while characterising unique *Cryptosporidium* genes from different countries [63]. However, a few genes identified here have been associated with increased virulence, host-pathogen interactions, or stress adaptation in parasites [64–67]. Further studies are necessary to assess the evolutionary and clinical significance of these genes, and others, in Australian *Cryptosporidium*.

Two isolates showed evidence of heterogeneity not previously detected using Sanger sequencing of the *SSU* and *gp60* loci [4]. The larger genome size recovered here for *C. meleagridis* appeared to be due to conflicting contig sequences, however the recovery of a single *gp60* and *SSU* from the assembly of this isolate suggests intra-specimen genomic heterogeneity showing nucleotide diversity or high allelic variation within this isolate. Whole genome amplification and assembly artefacts are also possible explanations. The low number (e.g. only four) of other *C. meleagridis* genome assemblies worldwide limits our ability to better evaluate intra-specific and intra-subtype diversity. The recovery of two distinct subtypes from AUSMDU00108955 suggests high allelic polymorphisms [68], presence of mixed populations [62, 69], or high rates of recombination in chromosome 6 at the *gp60* loci [61, 63]. High diversity of *gp60* sequences within the same host have been reported from New Zealand [68] and mixed population are often detected [62, 69]. Frequent and high rates of recombination between different *gp60* subtypes have been shown for *C. parvum* and *C. hominis* [61–63] with intra-specimen heterogeneity reported in the trinucleotide region [61]. Advancement in oocyst *in vitro* culturing and single-oocyst genomics [70, 71] could facilitate the resolution of such mixed populations in future studies.

The phylogenomic relationships of the Australian isolates are broadly consistent with previously described lineage structure within *C. hominis* and *C. parvum*. The phylogenetic relationships within *C. hominis hominis* do not appear to solely reflect geography nor subtype identity [8, 28, 62]. Additional genomes from poorly represented continent, countries, and subtypes is needed to further characterise the relationships within this subspecies and assess the similarity of isolates based on other genomic, epidemiological or phenotypic factors. Australian genomes belonging to the subspecies *C. hominis aquapotensis* would further inform the relationship between Australian and New Zealand isolates in comparison to North American [72].

The position of the Australian *C. parvum* placed in an intermediate clade between European and North American isolates within the *C. parvum parvum* subspecies would require additional genomes from Africa, Asia and Oceania to confirm the geographical signal within isolates. The Australian *C. parvum* isolate clustered within the Western lineage which has previously been associated with historical intercontinental livestock movement [19]. The position of the Australian isolate within the *C. p. parvum* subspecies is consistent with the broad host range of subtype IIaA20G3R1, which has been identified in livestock (cattle and alpacas) and wild deer in Australia [73–75]. The availability of more *Cryptosporidium* genomes from other isolates in the clinical and veterinary context is necessary to improve our understanding of genetic diversity and evolutionary pressures for this parasite [21, 41, 76]. A limitation of this study is that the phylogenetic relationships are based on a low number of Australian isolates especially for *C. parvum*. Additional clinical Australian isolates would help to validate these intra-specific relationships and improve our understanding of the population structure and phylogeography of *Cryptosporidium* in Australia. Genome assemblies of unique Australian *Cryptosporidium* species such as *C. macropodum, C.* sp. koala genotype, possum genotype, and the Australian mink genotype, among many others [4, 77, 78], would further elucidate the evolutionary relationships within *Cryptosporidium* in Australia. These genomes may provide insights into patterns of host adaptation, host movement and geographic diversification.

This study provides the necessary resources to enable genomic surveillance of cryptosporidiosis in Australia by presenting two locally-relevant reference genomes along with laboratory and bioinformatics workflows for clinical samples [41]. The routine genomic surveillance of clinical samples is still in its early days and has shown high genomic diversity within the traditional *gp60* subtypes and subfamilies [8, 19, 28, 61]. The implementation of routine molecular surveillance in the USA has helped public health authorities elucidate chains of outbreak transmission and epidemiology [15, 79]. Recent developments in clinical genomic surveillance has focused on the use of RNA bait approaches which offer direct capture from preserved stool samples but have limited recovery ability across the taxonomic range of *Cryptosporidium* species infecting humans [16, 80–82]. An advantage of the IMS approach is its ability to recover a wide range of species (usually from fresh faecal material) at lower cost and with less labour than RNA bait protocols [83]. In Australia, positive clinical faecal samples from diagnostics providers are kept fresh during diagnostics and referral to reference diagnostic laboratories, providing a practical approach for genomic surveillance. However, sample processing protocols in other countries could limit its application more generally. The combination of genomic surveillance in the clinical sector with surveillance in the veterinary and environmental sector offers a promising opportunity to investigate the transmission dynamics of cryptosporidiosis in humans and animals in Australia [4, 84, 85].

## CONCLUSION

This study provides reference genomes for *C. hominis* IaA12R3 and *C. parvum* IIaA20G3R1 and a draft genome assembly for *C. meleagridis* IIIgA31, representing *gp60* subtypes for which genomic assembly were previously unavailable, together with additional genomic data from nine Australian *C. hominis* isolates. The Australian genomes showed high levels of sequence similarity, orthology and synteny relative to other reference genomes. This study provides the genomic resources, bioinformatic workflows, and *Cryptosporidium* oocyst and DNA isolation protocols required to implement genomic surveillance of cryptosporidiosis in Australia across the clinical sector and beyond.

## Abbreviations

MDU PHL: Microbiological Diagnostics Unit Public Health Laboratory
ONT: Oxford Nanopore Technology
WGS: Whole genome sequencing
IMS: immunomagnetic separation
SNP: single nucleotide polymorphism

All supporting data, code and protocols have been provided within the article or through supplementary data files; Supplementary methods and two supplementary table are available with the online version of this article.

## Ethical approval

Ethical approval to conduct the study was received from the University of Melbourne Human Research Ethics Committee (approval number 2025-30320-66468-5).

## Acknowledgements

We thank the various pathology diagnostics service providers for sending samples to MDU PHL in 2024 and 2025. We thank the laboratory technical staff at MDU PHL for samples reception, cataloguing and storage. We thank Jeanne Arona and Rangimarie Whatley (BioPoint Pty) for technical assistance with IMS protocol.

## Author contributions

MB methodology, formal analysis, investigation and writing original draft. SS methodology and validation. KM and MS conceptualization, methodology, validation, resources. THH and LJ methodology, resources, investigation, data curation. NS resources, writing review & editing. JL data curation, software. AVK methodology, writing review and editing. RW methodology, software, validation, formal analysis, review. TS methodology, software, formal analysis, writing review and editing. BPH and RBG funding acquisition, supervision, conceptualization, resources, writing review & editing.

