## Supplementary Figures for "Reference genomes for Australian *Cryptosporidium hominis* and *C. parvum* from humans"

### Slide 1
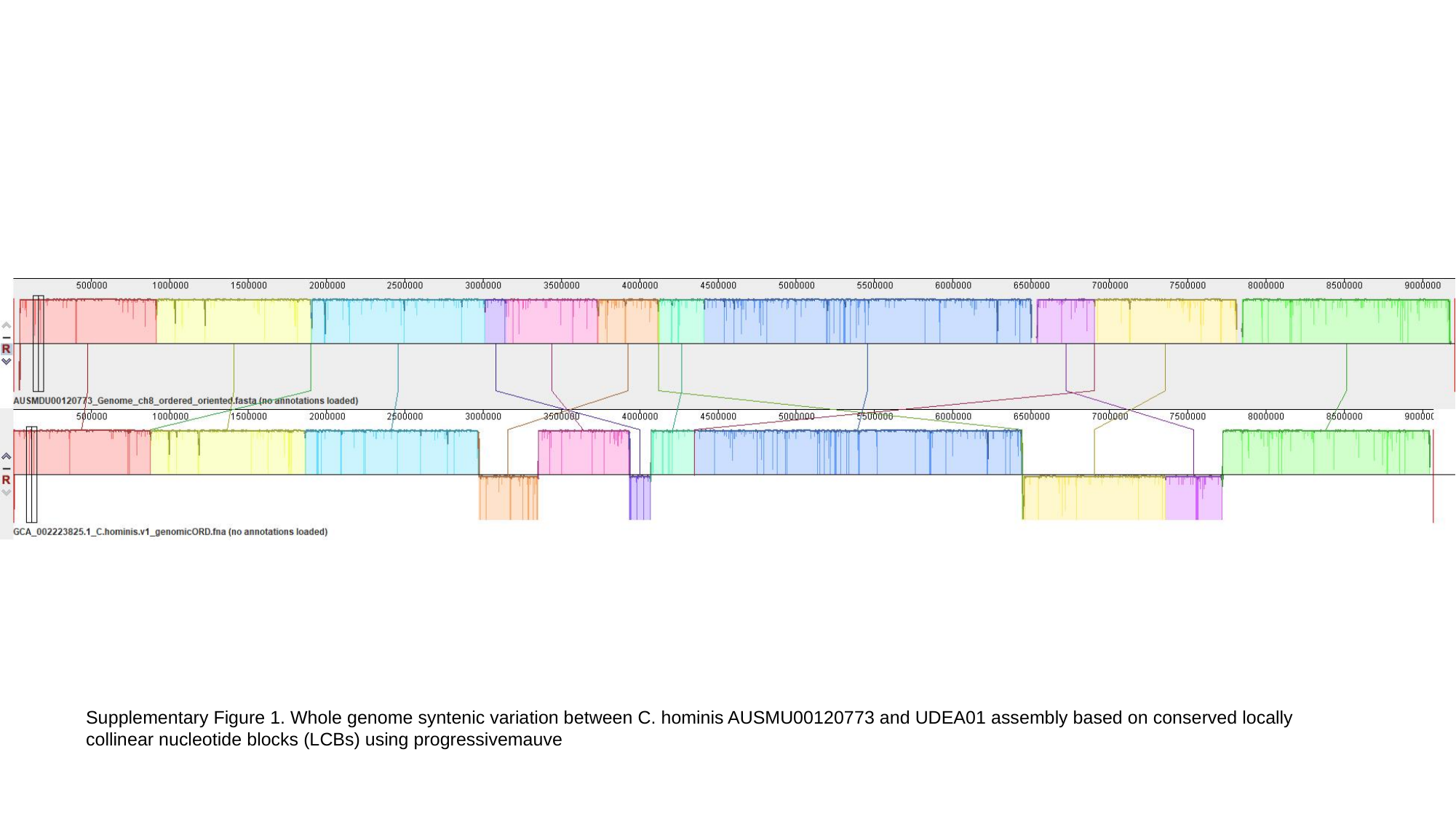

Supplementary Figure 1. Whole genome syntenic variation between C. hominis AUSMU00120773 and UDEA01 assembly based on conserved locally collinear nucleotide blocks (LCBs) using progressivemauve

### Slide 2
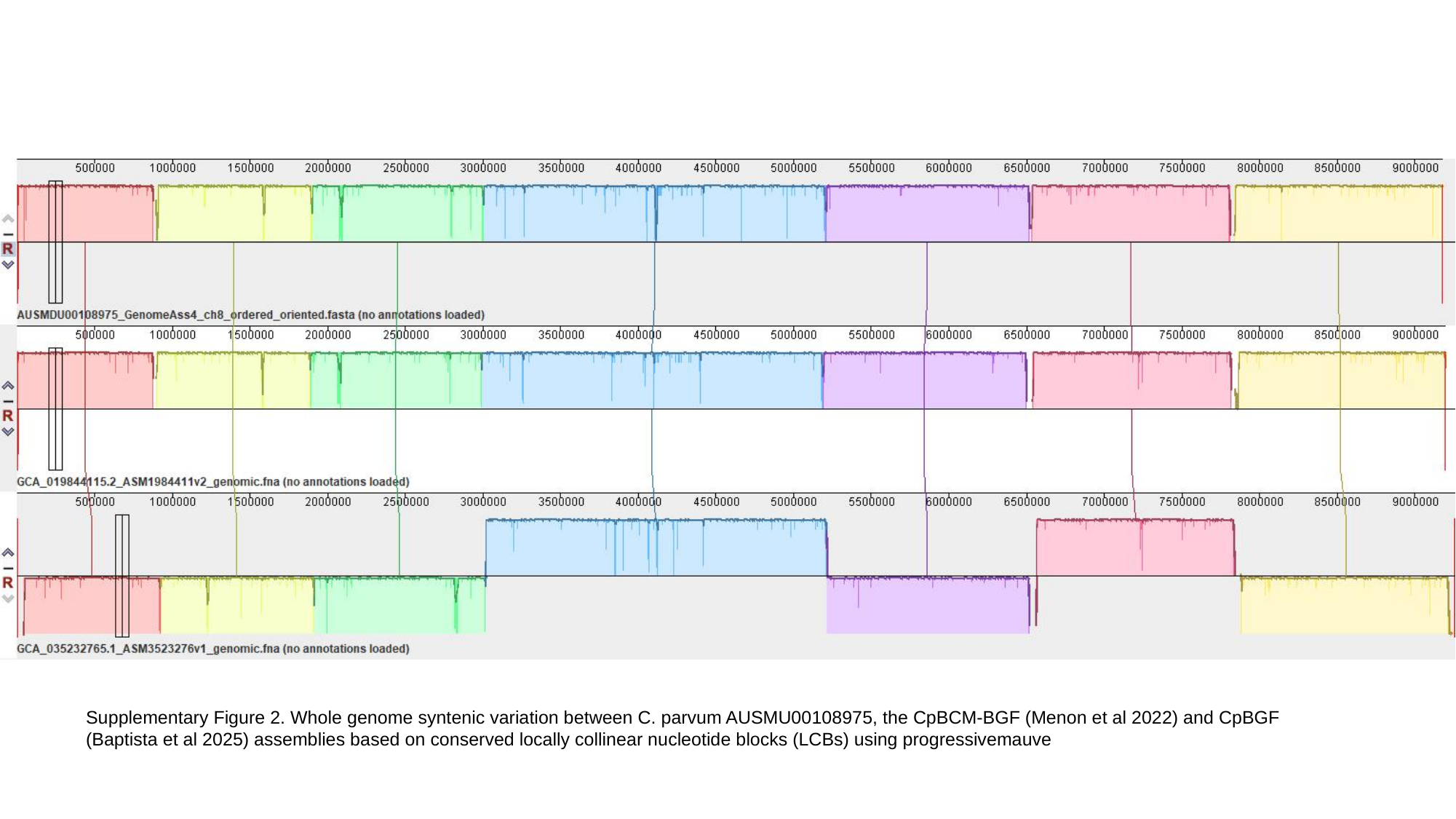

Supplementary Figure 2. Whole genome syntenic variation between C. parvum AUSMU00108975, the CpBCM-BGF (Menon et al 2022) and CpBGF (Baptista et al 2025) assemblies based on conserved locally collinear nucleotide blocks (LCBs) using progressivemauve

### Slide 3
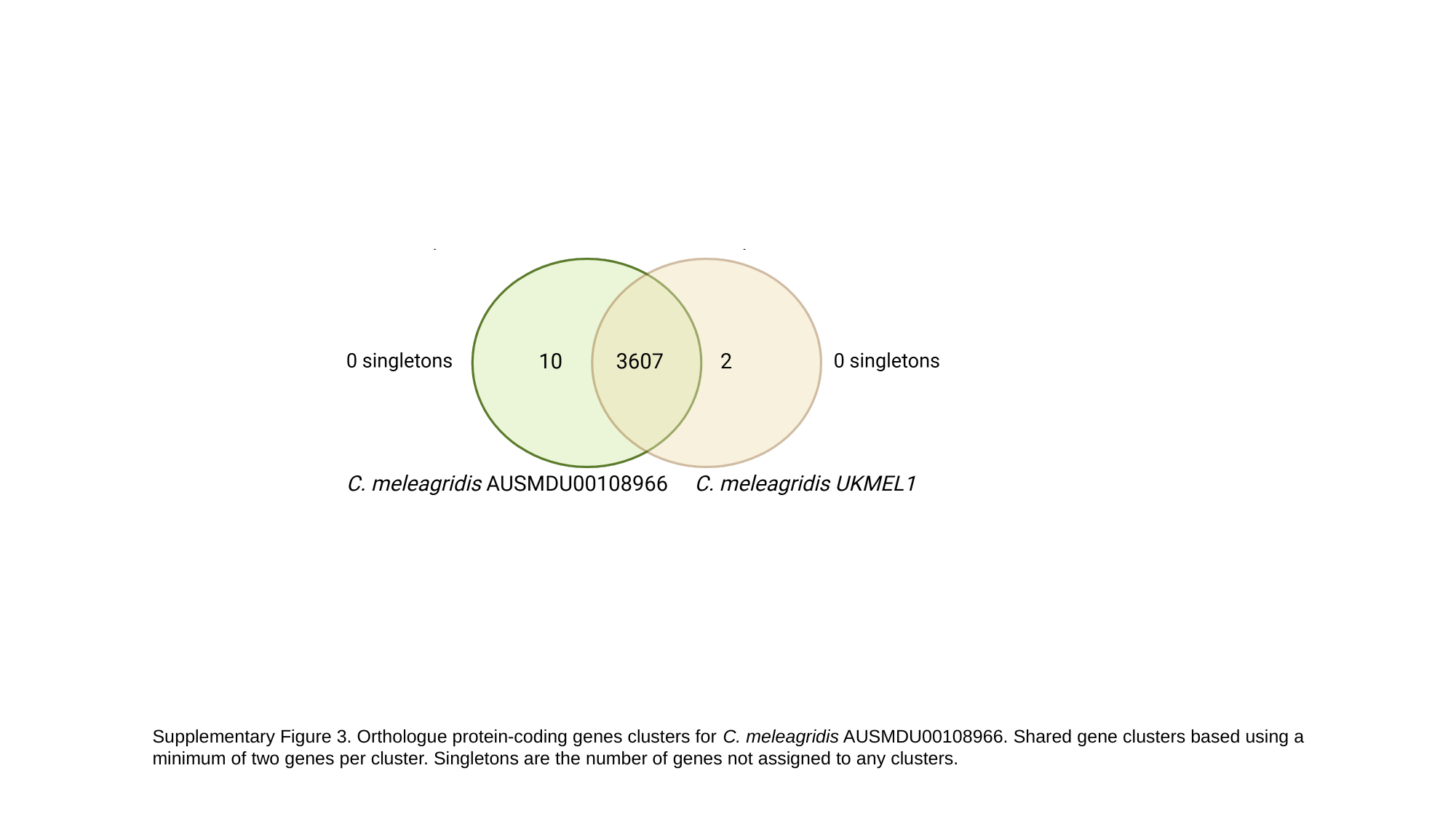

Supplementary Figure 3. Orthologue protein-coding genes clusters for C. meleagridis AUSMDU00108966. Shared gene clusters based using a minimum of two genes per cluster. Singletons are the number of genes not assigned to any clusters.

### Slide 4
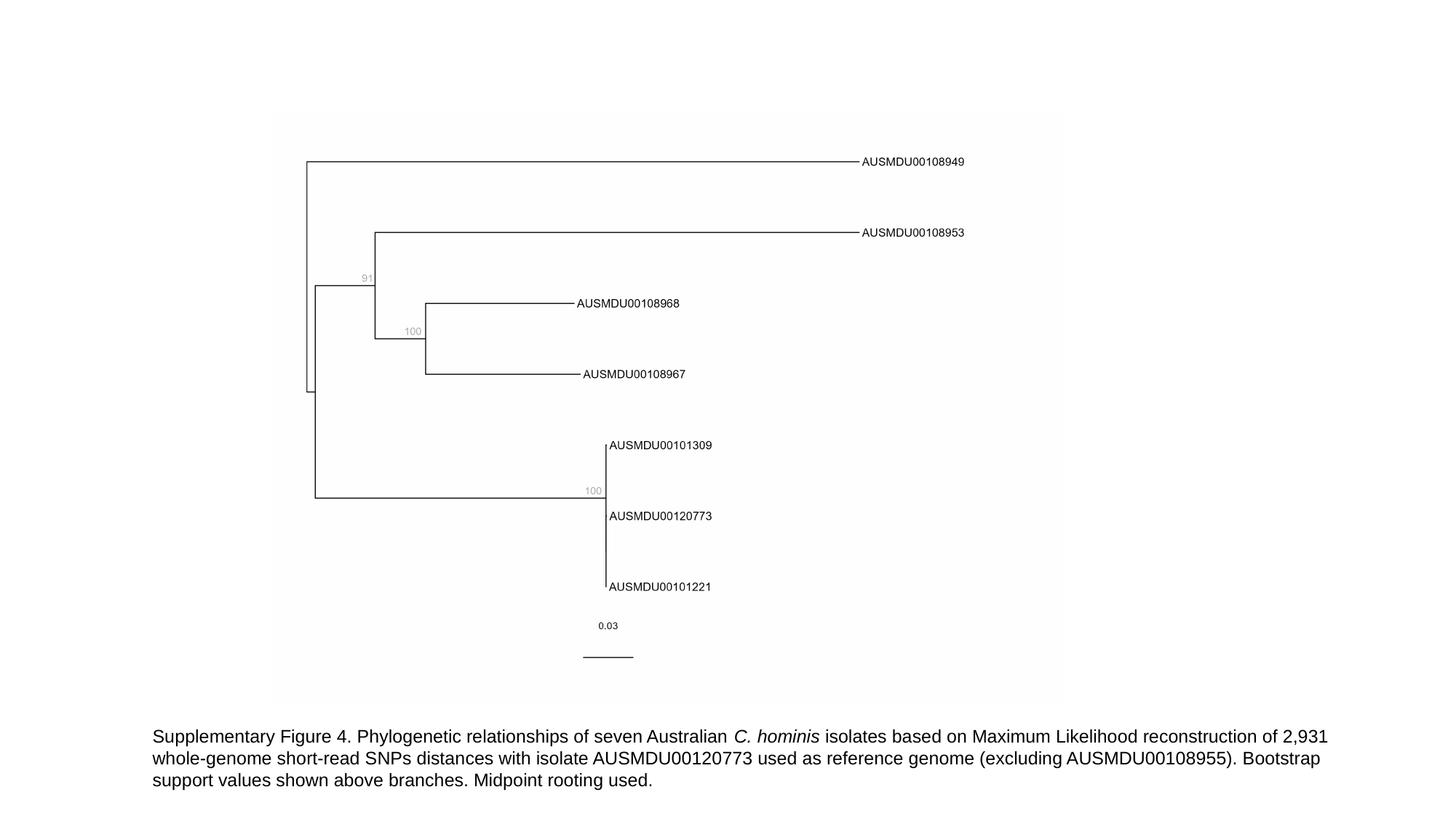

Supplementary Figure 4. Phylogenetic relationships of seven Australian C. hominis isolates based on Maximum Likelihood reconstruction of 2,931 whole-genome short-read SNPs distances with isolate AUSMDU00120773 used as reference genome (excluding AUSMDU00108955). Bootstrap support values shown above branches. Midpoint rooting used.
