## Supplementary Table 3 for "Reference genomes for Australian *Cryptosporidium hominis* and *C. parvum* from humans"

Supplementary Table 3. Genome assembly and annotation statistics for the draft *C. meleagridis* genomes (AUSMDU) compared to same species reference genome.

|  | *C. meleagridis* | |
| --- | --- | --- |
| Isolate | AUSMDU00108966 | TU1867 |
| Number of assembled chromosomes | 0 | 8 |
| Number of contigs | 113 | 13 |
| Number of telomeres | 0 | 1 |
| Genome size | 9,370,719 | 9,178,484 |
| Genome coverage | 157x | 361x |
| Longest contig (bp) | 515,062 | 1,365,597 |
| N50 | 216,879 | 1,105,563 |
| N90 | 64,986 | 880,501 |
| L50 | 14 | 4 |
| Number of N per 100 kbp | 0 | 0 |
| Overall GC % | 31.0 | 31.9 |
| Number of genes | 4,075 | 3,988 |
| Protein-coding genes / non-coding genes | 3,981 / 94 | 3,919 / 69 |
| Complete & single-copy gene (%, M; Missing, F; Fragmented) | 468 (97.2%, 10M, 4F) | 493 (98.4%, 5M, 3F) |
