## Supplementary Methods for "Reference genomes for Australian *Cryptosporidium hominis* and *C. parvum* from humans"

**Novel Australian *Cryptosporidium* genomes from previously uncharacterised subtypes of *C. hominis*, *C. parvum* and *C. meleagridis***

Supplementary Methods

**Modified Immunomagnetic separation (IMS) protocol**

Approximately 1g of faecal material was added to a 15 mL tube containing 1 mL of Buffer B, 1 mL of Buffer A, and 5-7 mL of purified water (equilibrated to room temperature without crystallization). Vortex the sample at maximum speed for 60 seconds or until the sample is homogeneously distributed and has the liquid consistency of water or slightly more viscous. Transfer 100 ul of CryptoBeads® and invert tube 10x to mix well. Insert the tube on a vertical rotator and rotate at 20 rpm for 1 hour. Attach the tube to magnet and rotate at 10 rpm for 5 minutes. Discard supernatant while holding the magnet up so that the pelleted magnetic beads are not disturbed. Aspirate any leftover liquid. Remove the tube from the magnet and add 8 mL of PBS 1X, invert the tube 10x to mix well. Place the tube in the magnet and rotate at 10 rpm for 5 minutes. While on magnet, discard the supernatant. Perform another PBS wash if sample still contains a large amount of faecal material. Add 1.2 mL of PBS and vortex gently. Transfer liquid to 2 mL low-protein binding tube, attach to a magnet and rotate at 10 rpm for 3 minutes. While on magnet, discard the supernatant ensuring all the liquid is removed including from inside the lid. Remove from magnet, add 100 ul of 0.1M HCl and vortex at maximum speed for 60 seconds. Incubate at room temperature for 10 minutes, followed by vortexing at maximum speed for 60 seconds. Place on the magnetized dissociation bloc for 2 minutes. While on the magnetized dissociation bloc, transfer the supernatant to a new low-protein binding tube and add 10 ul of 1M NaOH to the purified oocysts. Repeat dissociation step another time to obtain the maximum number of oocysts attached the to the beads.

**Modified DNA extraction protocol**

DNA extraction protocol was similar to Babineau et al 2025 JCM and used a DNA Lysis Buffer consisting of 10% SDS, 1M Tris, 1M NaCl, 0.5M EDTA. Modified steps include a vortexing step between the seven freeze-thaw cycles, the use of Chloroform:Isoamyl 24:1 (Thermo Fisher Scientific, USA) for the second purification step, DNA precipitation in 100% isopropanol overnight at room temperature, the removal of all post freeze-thaw vortexing steps. Final elution used 25 ul of purified water.

**Modified T7 debranching protocol**

Amplified DNA was purified using AMPure XP (Agencourt #A63880) beads following the Oxford Nanopore protocol v114 for ligation and sequencing of genomic DNA with final elution of 25 ul. The whole volume was then combined with 3 ul of NEB Buffer2 and 3 ul of T7 endonuclease I (New England Biolabs # M0302S, UK) and incubated at 37°C for a minimum of 60 minutes. Then 90 ul of custom Buffer, 20 ul of TE Buffer was added to the DNA and cleaned using the AMPure XP bead protocol and eluted in 49 ul.
